## Supplementary figures for "Carafe enables high quality *in silico* spectral library generation for data-independent acquisition proteomics"

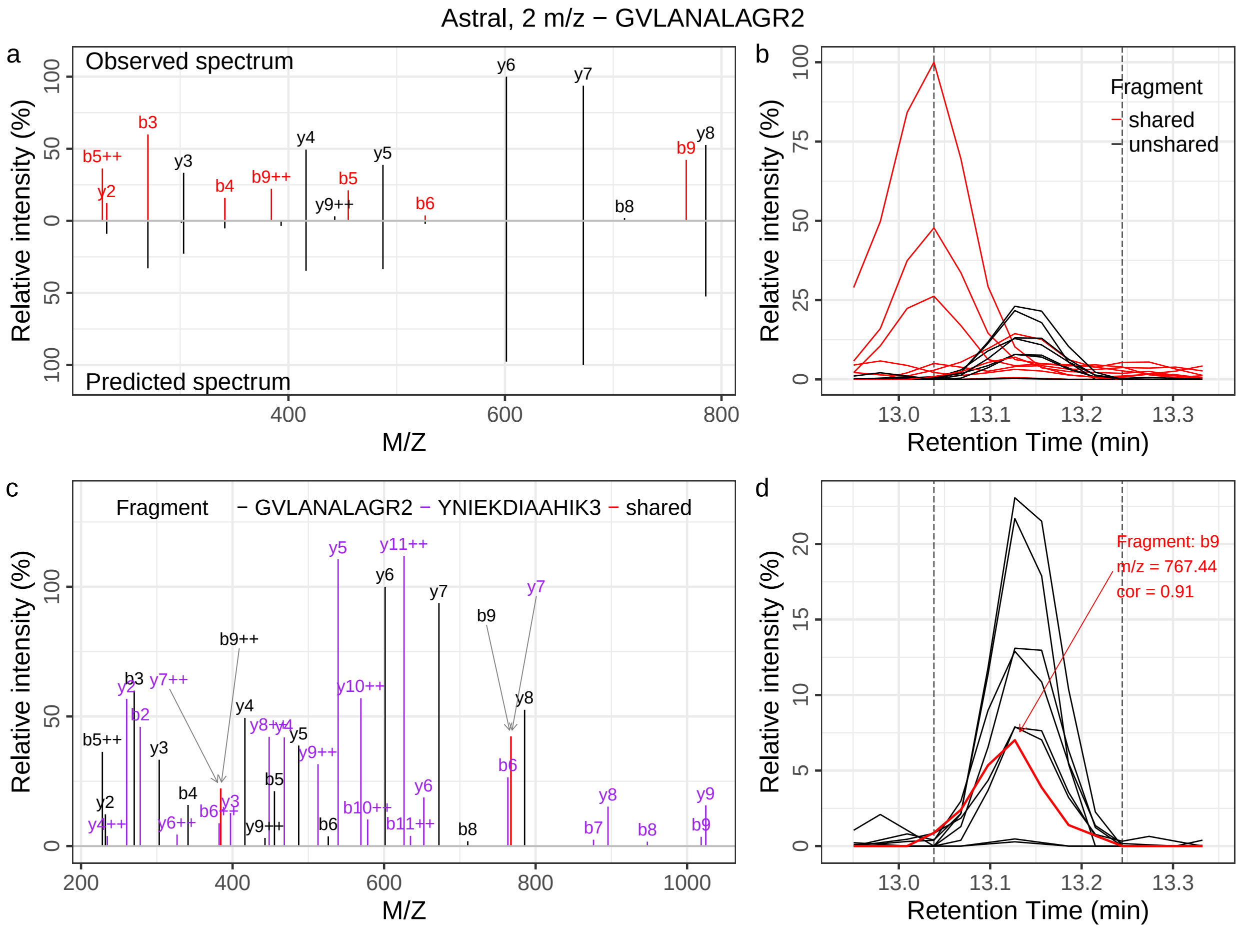


Figure S1: **A shared peak detection example using the two-pronged approach in Carafe. (a)** The top panel shows the observed fragment ions for peptide GVLANALAGR with precursor charge 2+. The fragments highlighted as red color are shared fragments determined using Carafe. The bottom panel shows the predicted fragment ion intensity values using the pretrained AlphaPeptDeep model. **(b)** Extracted ion chromatograms (XICs) of all observed fragment ions matched to peptide GVLANALAGR. The fragments highlighted as red color are shared fragments determined using Carafe. The dashed lines indicate the peak boundaries. **(c)** A different peptide YNIEKDIAAHIK with precursor charge 3+ was detected from the same MS2 spectrum that was used to detect the peptide GVLANALAGR showing in **(a)**. Only observed fragment ions matched to the two peptides were shown. The fragment ions annotated to both peptides are highlighted as red color. **(d)** XICs of the observed fragment ions matched to peptide GVLANALAGR which were determined as unshared peaks using the peptide-centric method by Carafe. The fragment highlighted as red color (b9) was only determined as a shared peak by the spectrum-centric method by Carafe. This fragment was not determined as a shared peak using the peptide-centric method due to its high correlation (Pearson correlation: 0.91) to the best fragment ion XIC of the peptide.

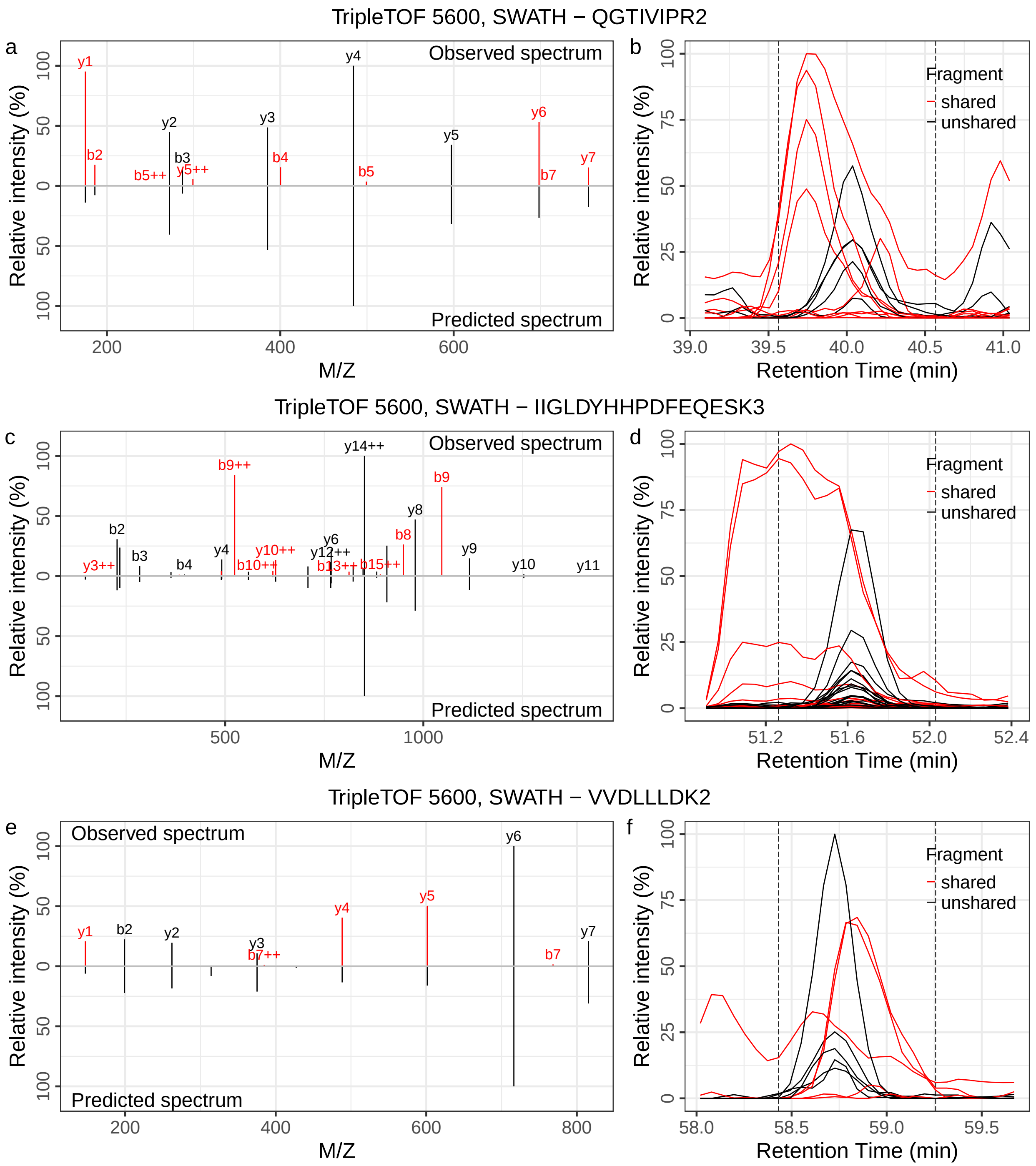


Figure S2: **Examples of shared peak detection by Carafe on the TripleTOF5600 dataset. (a)**, **(c)** and **(e)** show the observed fragment ions and predicted fragment ion intensity values for the three peptides. The fragments highlighted as red color are shared fragments determined using Carafe. **(b)**, **(d)** and **(f)** show the XICs of all observed fragment ions matched to the corresponding peptides. The fragments highlighted as red color are shared fragments determined using Carafe. The dashed lines indicate the peak boundaries.

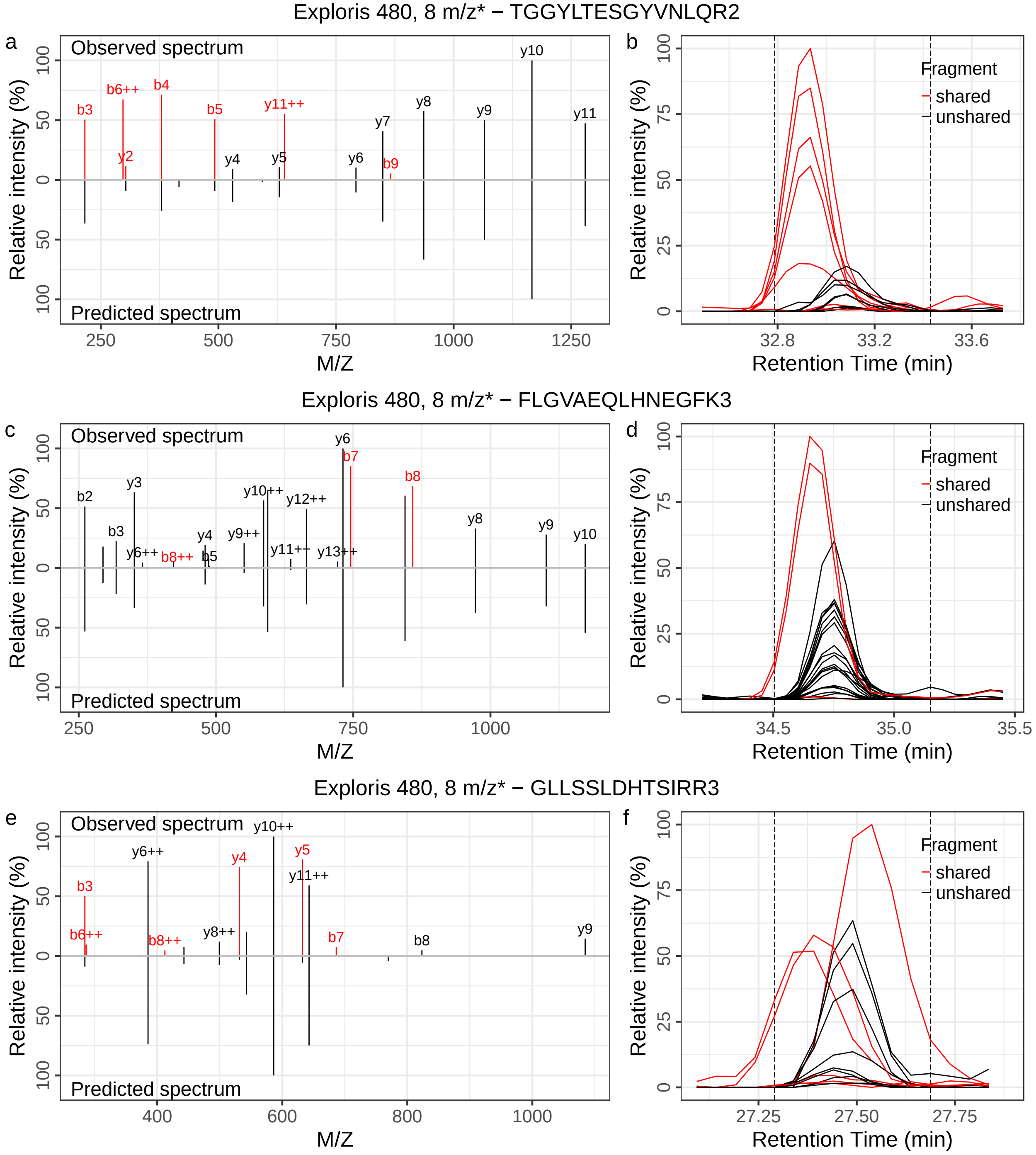


Figure S3: **Examples of shared peak detection by Carafe on the Exploris 480 metaproteome dataset.** **(a)**, **(c)** and **(e)** show the observed fragment ions and predicted fragment ion intensity values for the three peptides. The fragments highlighted as red color are shared fragments determined using Carafe. **(b)**, **(d)** and **(f)** show the XICs of all observed fragment ions matched to the corresponding peptides. The fragments highlighted as red color are shared fragments determined using Carafe. The dashed lines indicate the peak boundaries.

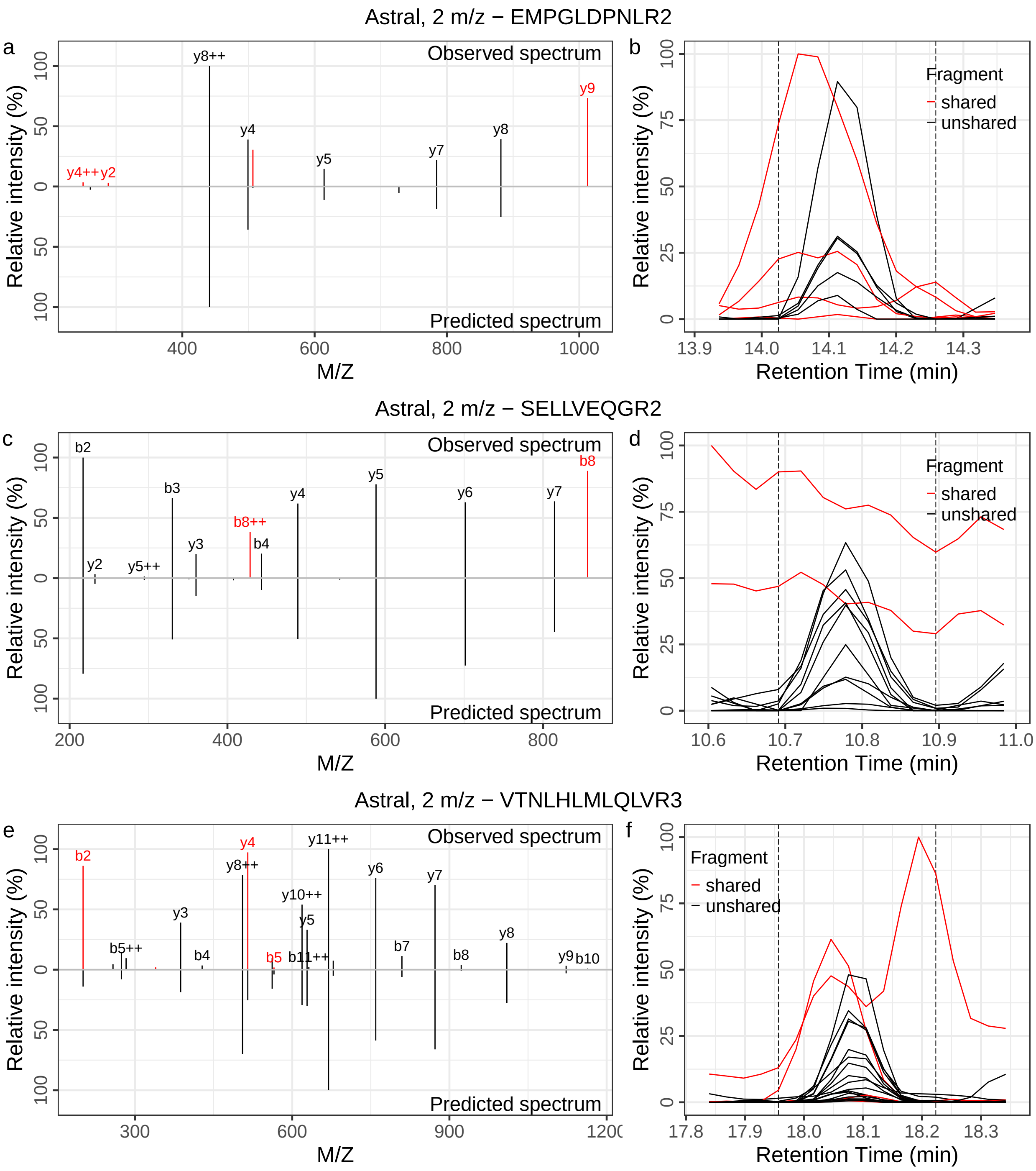


Figure S4: **Examples of shared peak detection by Carafe on the Astral dataset. (a)**, **(c)** and **(e)** show the observed fragment ions and predicted fragment ion intensity values for the three peptides. The fragments highlighted as red color are shared fragments determined using Carafe. **(b)**, **(d)** and **(f)** show the XICs of all observed fragment ions matched to the corresponding peptides. The fragments highlighted as red color are shared fragments determined using Carafe. The dashed lines indicate the peak boundaries.

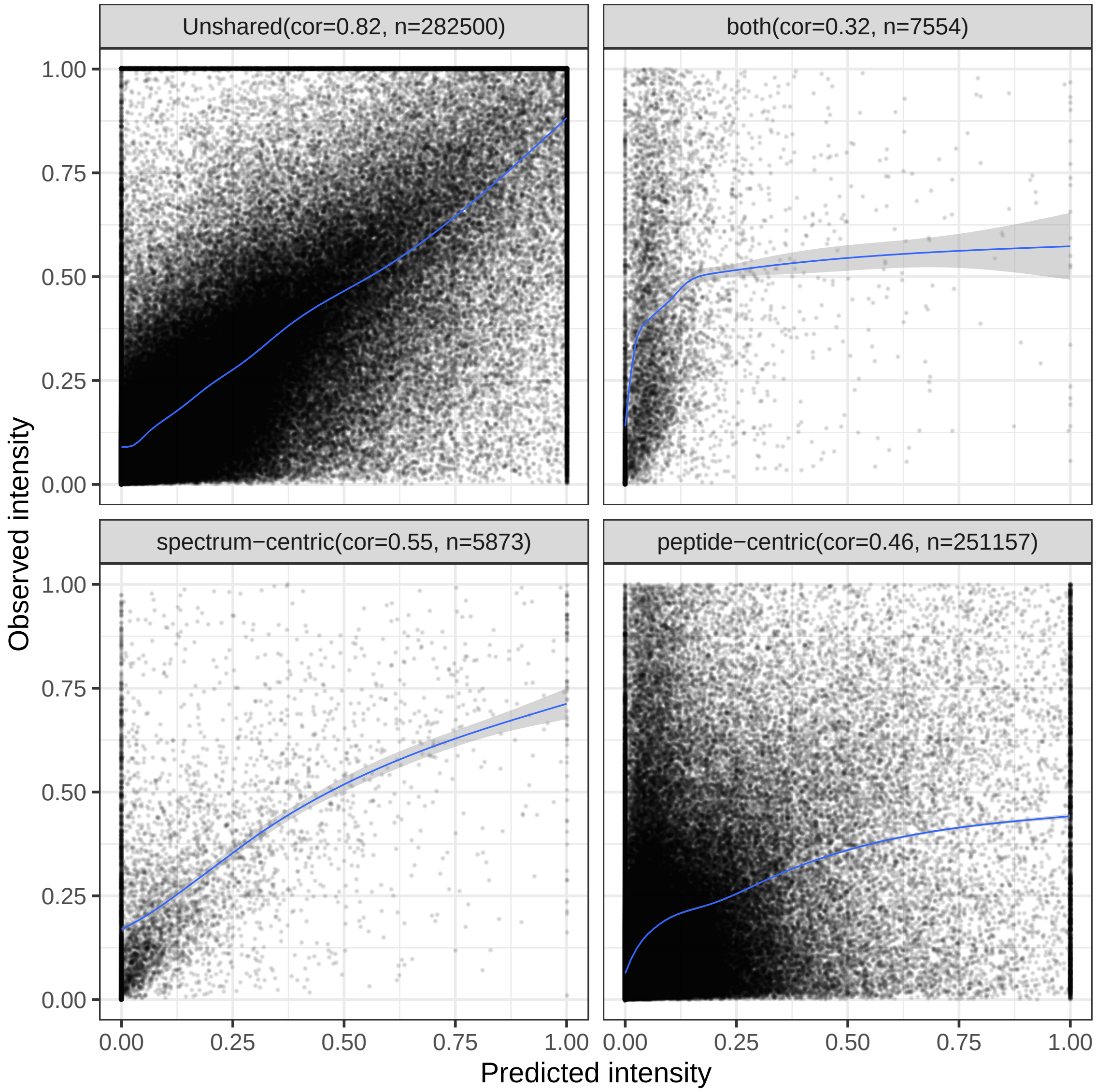


Figure S5: **The performance of the shared peak detection methods.** The detected peaks were divided into four groups: detected as interfered by one method, the other, both or neither (unshared). Each panel plots the predicted versus observed peak intensities, and the blue lines represent a smooth trend calculated using a loess method, highlighting the general pattern of correlation across the intensity range. The number of points and the Pearson correlation between observed and predicted fragment ion intensities is shown at the top of each panel.

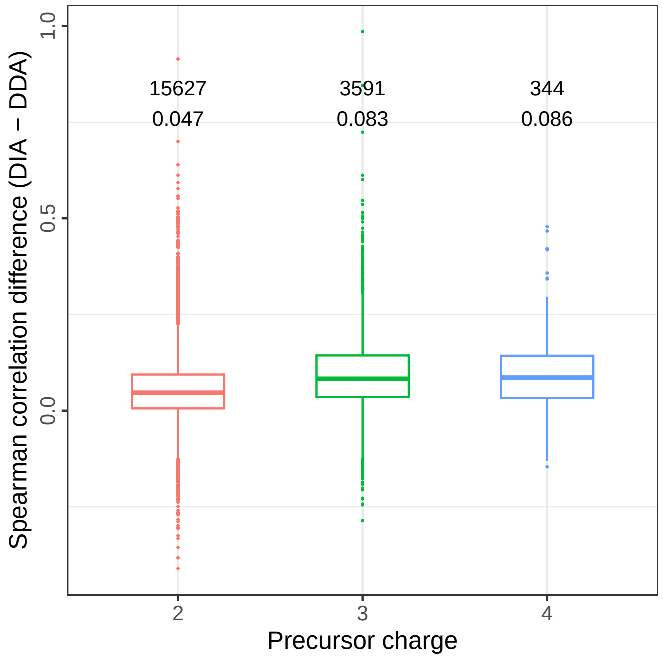


Figure S6: **Fragment ion intensity prediction improvement across precursor charge states.** Spearman correlation coefficients between predicted and observed fragment ion intensities were calculated for each peptide precursor using both the DIA fine-tuned model and the pretrained DDA model on the TripleTOF 5600 dataset. The y-axis shows the difference in Spearman correlation (fine-tuned minus pretrained) for each precursor charge state. The numbers in the first row in the plot are the number of peptide precursors in each precursor charge state group used for the analysis. The numbers in the second row are the median Spearman correlation difference values for each precursor charge state group. In the boxplot, centerline indicates the median, the lower and upper hinges correspond to the first and third quartiles, whiskers indicate the 1.5 interquartile range and points indicate outliers.

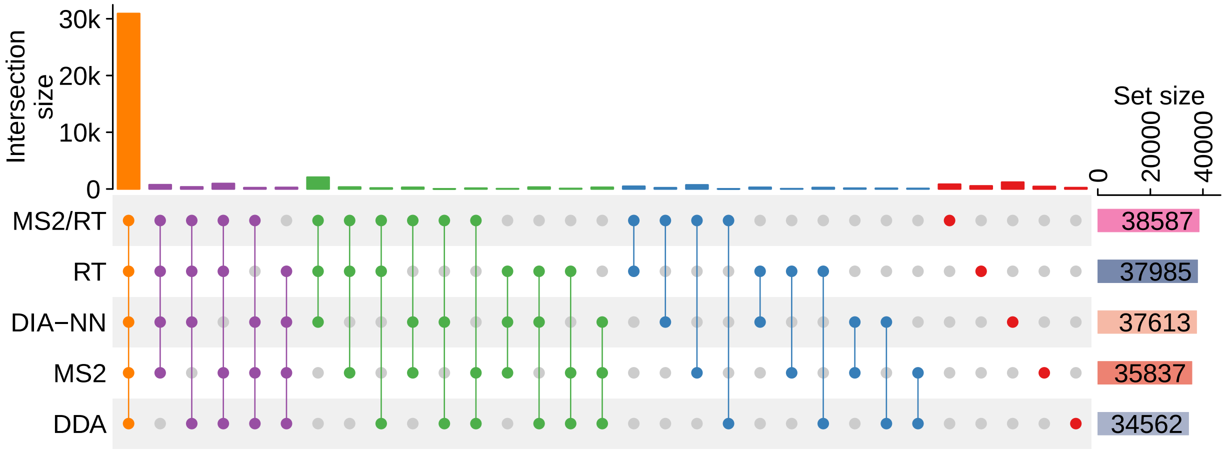


Figure S7: **Upset plot on the TripleTOF 5600 dataset.** The dots representing combination sets are colored according to the number of sets included.

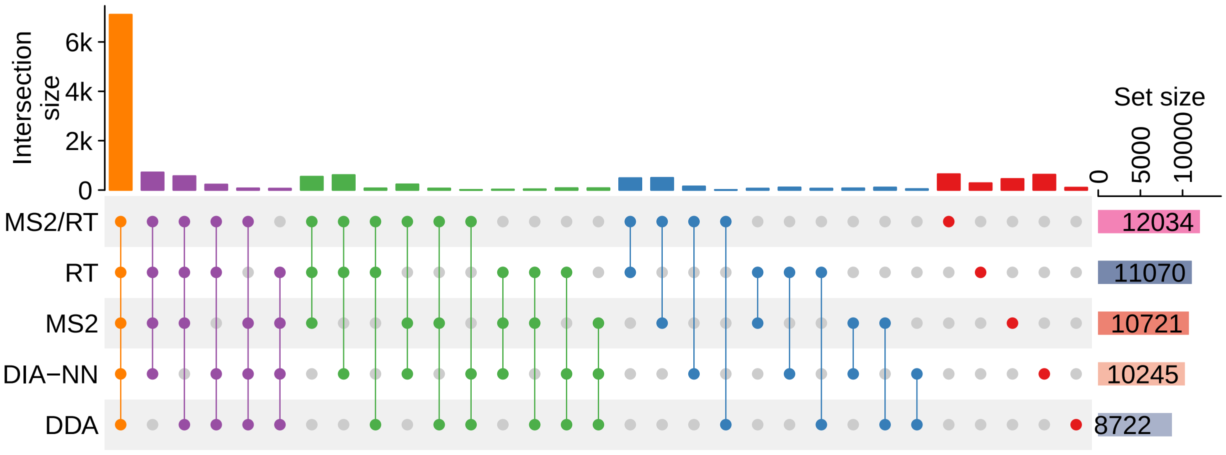


Figure S8: **Upset plot on the metaproteomics DIA dataset.** The dots representing combination sets are colored according to the number of sets included.

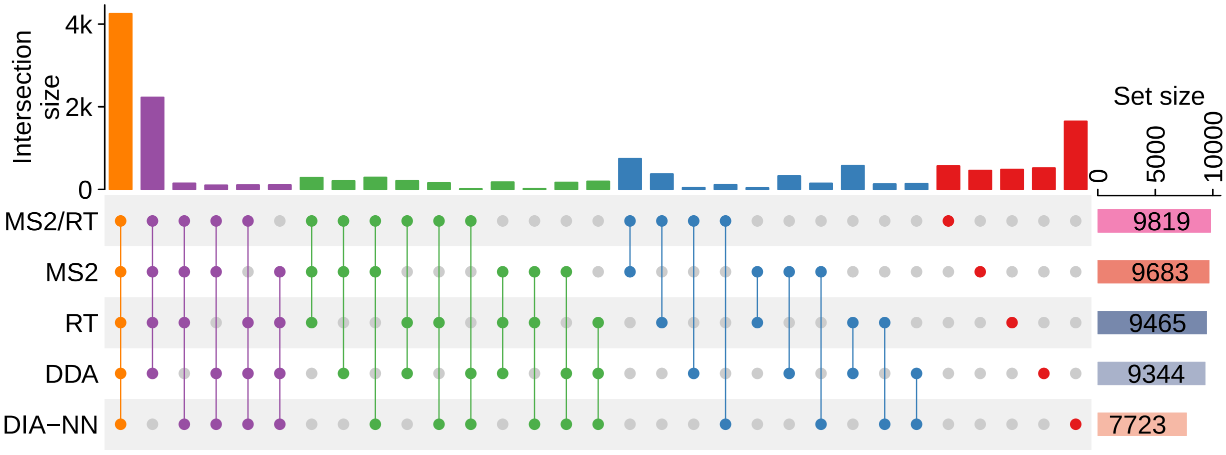


Figure S9: **Upset plot on the phosphoproteomics DIA dataset.** The dots representing combination sets are colored according to the number of sets included.

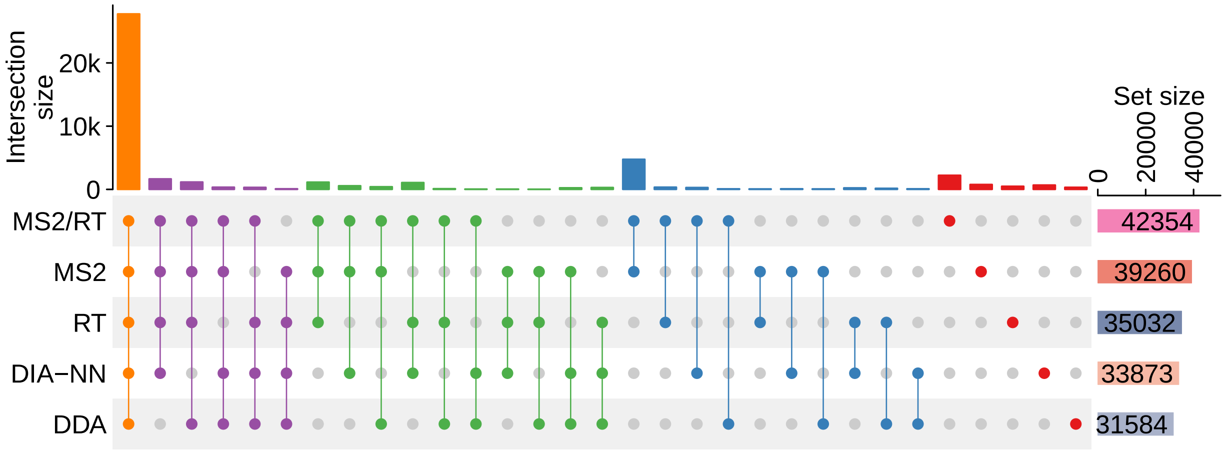


Figure S10: **Upset plot on the Lumos reCID dataset.** The dots representing combination sets are colored according to the number of sets included.

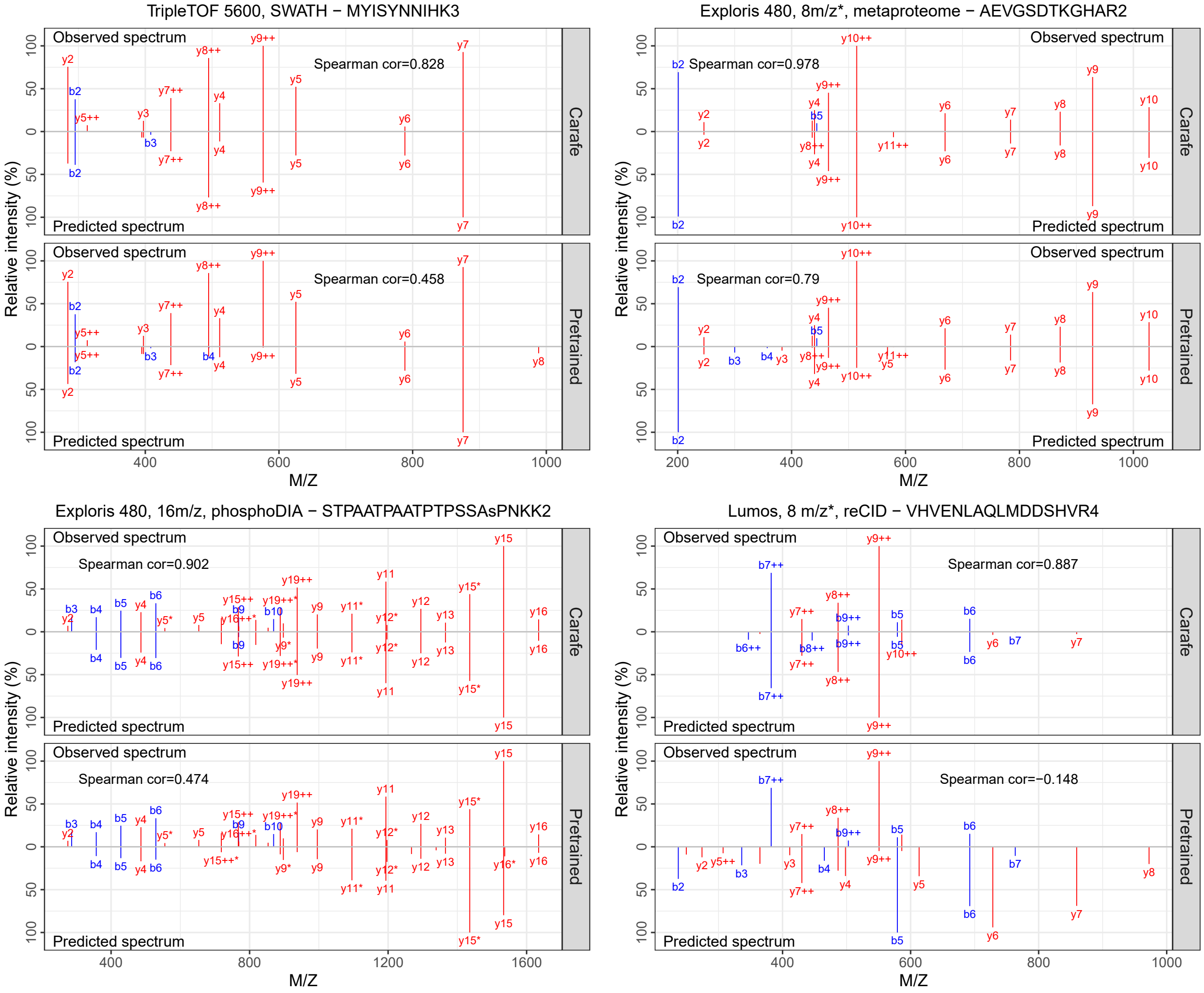


Figure S11: **Annotated spectra of four peptides detected using Carafe fine-tuned spectral libraries but not detected using the pretrained model (AlphaPeptDeep) derived spectral libraries.** For each subplot, the top panel shows the experimental spectrum while the bottom panel shows the predicted spectrum. Only observed fragment ions matched to the corresponding peptide were shown.

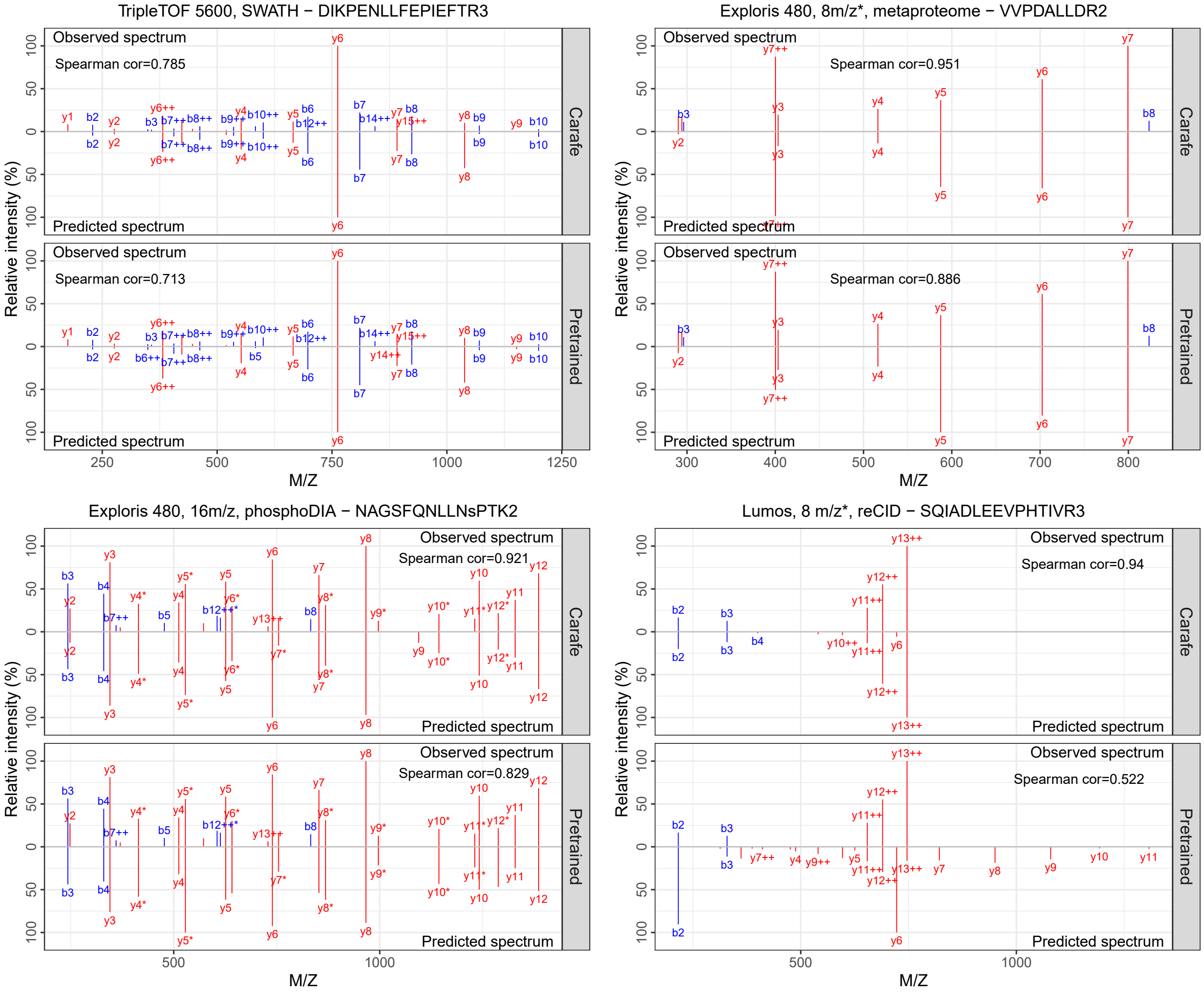


Figure S12: **Annotated spectra of four peptides detected using Carafe fine-tuned spectral libraries but not detected using the pretrained model (AlphaPeptDeep) derived spectral libraries.** For each subplot, the top panel shows the experimental spectrum while the bottom panel shows the predicted spectrum. Only observed fragment ions matched to the corresponding peptide were shown.

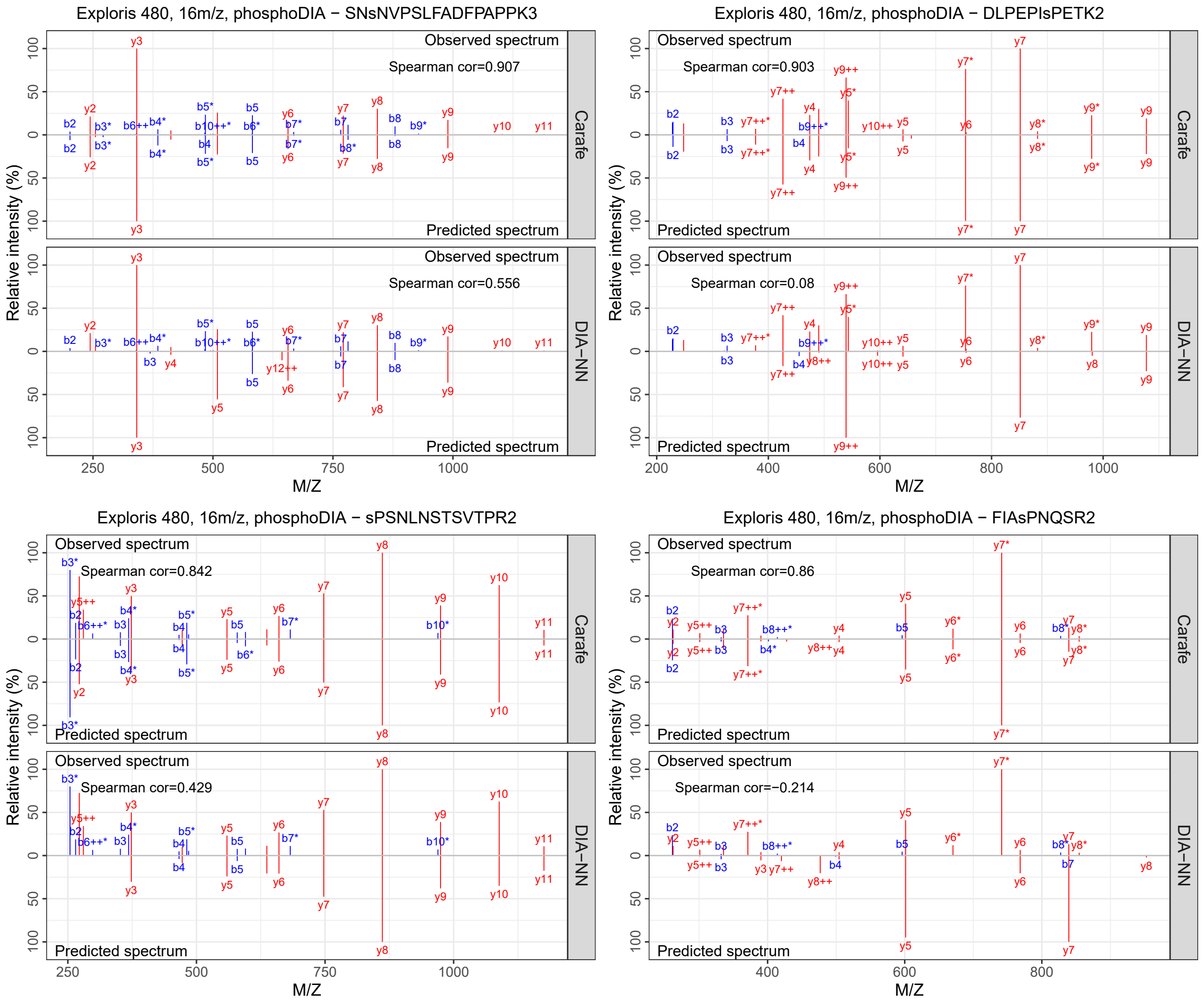


Figure S13: **Annotated spectra of four phosphopeptides detected using the Carafe fine-tuned spectral library but not detected using the DIA-NN generated spectral library.** For each subplot, the top panel shows the experimental spectrum while the bottom panel shows the predicted spectrum. Only observed fragment ions matched to the corresponding peptide were shown.

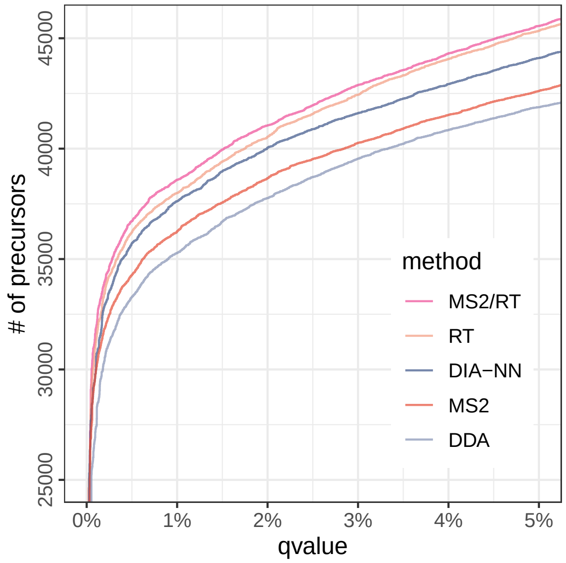


Figure S14: The number of precursors detected using DIA-NN with different spectral libraries as a function of qvalue threshold on the TripleTOF 5600 dataset. DDA: spectral library generated using the pretrained DDA model from AlphaPeptDeep; MS2: spectral library generated using a fine-tuned fragment ion intensity prediction model but the pretrained RT model from AlphaPeptDeep; RT: spectral library generated using a fine-tuned RT prediction model but the pretrained fragment ion intensity model from AlphaPeptDeep; MS2/RT: spectral library generated using fine-tuned fragment ion intensity and RT prediction models; DIA-NN: spectral library generated using DIA-NN.

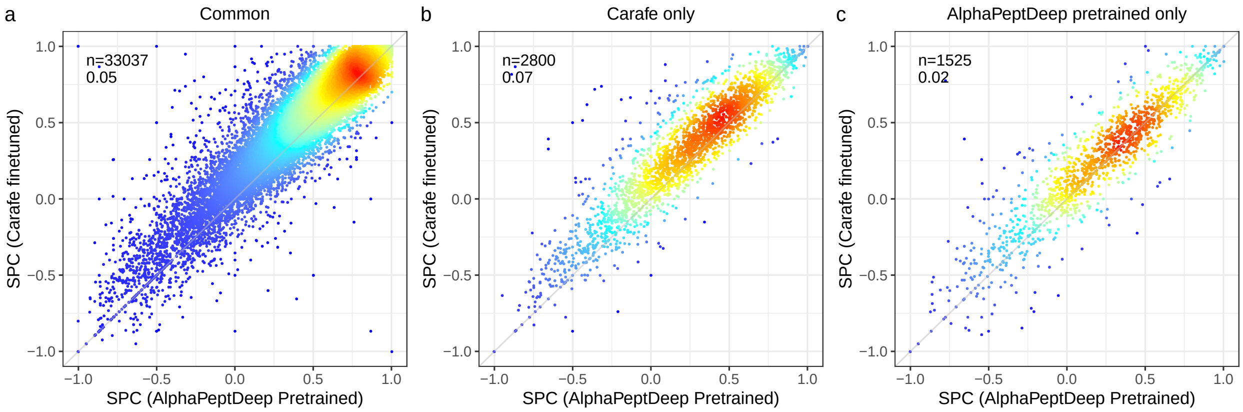


Figure S15: **Spearman correlation improvement analysis for precursors detected on the TripleTOF 5600 yeast dataset.** **(a)** Spearman correlation comparison for the precursors detected by both the Carafe fine-tuned spectral library (only fine-tuned fragment ion intensity prediction model, referred as "MS2" in Figure 4) and the spectral library generated from the AlphaPeptDeep pretrained model (referred as "DDA" in Figure 4). **(b)** Same as **(a)** but for the precursors only detected using the "MS2" library. **(c)** Same as **(a)** but for the precursors only detected using the "DDA" library. SPC: Spearman correlation. In each plot, $n$ is the number of precursors used in analysis. The value in the second row is median Spearman correlation improvement comparing the predictions using Carafe fine-tuned model with those using the AlphaPeptDeep pretrained model. Only peaks determined to be non-interfered were used in the correlation calculation.

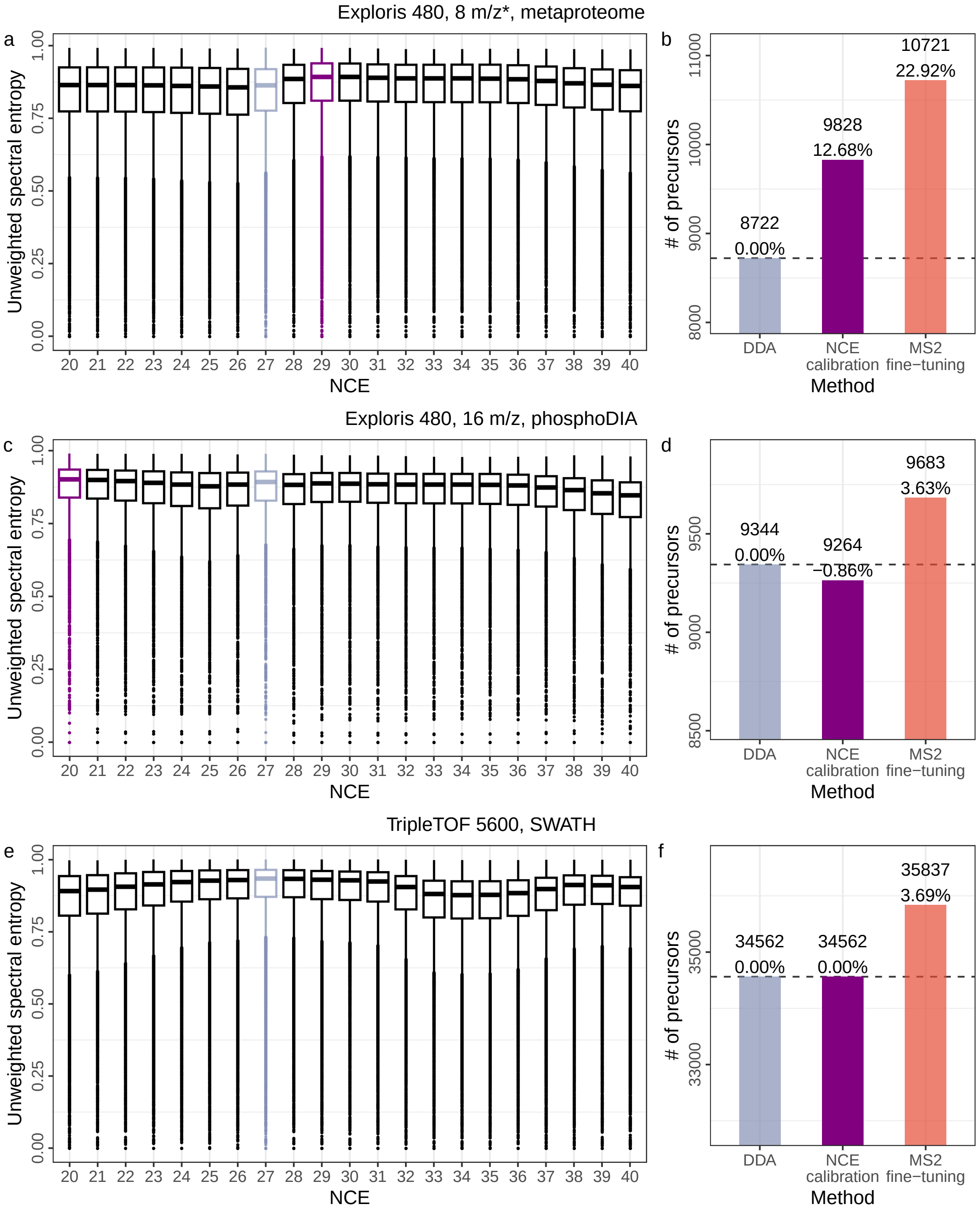


Figure S16: **NCE calibration performance evaluation**. **(a)**, **(c)** and **(e)** X axis shows the NCE values considered during the optimization. Y axis shows the unweighted spectral entropy values. The best NCE was determined by highest median entropy value and was highlighted with purple color. The NCE value used for generating a dataset from a Thermo instrument was highlighted as light blue. The NCE value used for the TripleTOF 5600 dataset by Carafe fine-tuning was highlighted as light blue. **(b)**, **(d)** and **(f)** show the number of precursors detected passed 1% FDR using DIA-NN with different spectral libraries. DDA: a spectral library generated using the pretrained DDA model from AlphaPeptDeep with the NCE used for generating the dataset or a default of 27 for the TripleTOF 5600 dataset. NCE calibration: a spectral library generated using the pretrained DDA model with the optimized NCE. MS2 fine-tuning: a spectral library generated using Carafe with only fragment ion intensity prediction fine-tuned. For boxplots, centerline indicates the median, the lower and upper hinges correspond to the first and third quartiles, whiskers indicate the 1.5 interquartile range and points indicate outliers.

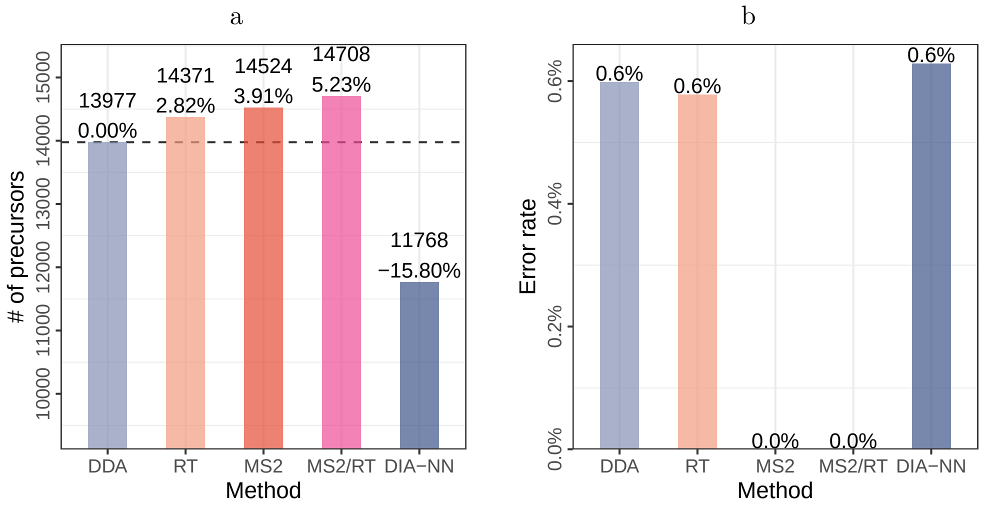


Figure S17: **Phosphopeptide detection error rate evaluation.** **(a)** Phosphopeptide precursors detected using different spectral libraries. **(b)** False localization rate evaluation using synthetic phosphopeptides detected with different spectral libraries.
